# DNSS2: improved *ab initio* protein secondary structure prediction using advanced deep learning architectures

**DOI:** 10.1101/639021

**Authors:** Jie Hou, Zhiye Guo, Jianlin Cheng

**Affiliations:** Department of Electrical Engineering and Computer Science, University of Missouri, Columbia, Missouri, 65211, USA

## Abstract

**Motivation:** Accurate prediction of protein secondary structure (alpha-helix, beta-strand and coil) is a crucial step for protein inter-residue contact prediction and *ab initio* tertiary structure prediction. In a previous study, we developed a deep belief network-based protein secondary structure method (DNSS1) and successfully advanced the prediction accuracy beyond 80%. In this work, we developed multiple advanced deep learning architectures (DNSS2) to further improve secondary structure prediction.

**Results:** The major improvements over the DNSS1 method include (i) designing and integrating six advanced one-dimensional deep convolutional/recurrent/residual/memory/fractal/inception networks to predict secondary structure, and (ii) using more sensitive profile features inferred from Hidden Markov model (HMM) and multiple sequence alignment (MSA). Most of the deep learning architectures are novel for protein secondary structure prediction. DNSS2 was systematically benchmarked on two independent test datasets with eight state-of-art tools and consistently ranked as one of the best methods. Particularly, DNSS2 was tested on the 82 protein targets of 2018 CASP13 experiment and achieved the best Q3 score of 83.74% and SOV score of 72.46%. DNSS2 is freely available at: https://github.com/multicom-toolbox/DNSS2.

## 1 Introduction

Three major types of protein secondary structure are alpha-helix (H), beta-strand (E) and coil state (C) (Pauling, et al., 1951), each of which represents the local structure state of an amino acid in a folded polypeptide chain. The predicted information of protein secondary structure is useful for many applications in computational biology, such as protein residue-residue contact prediction (Adhikari, et al., 2017; Michel, et al., 2018; Wang, et al., 2017), protein folding (Hou, et al., 2017; Jones, et al., 1999; Myers and Oas, 2001), *ab-initio* protein structure modeling (Adhikari and Cheng, 2018; Rohl, et al., 2004; Roy, et al., 2010) and protein model quality assessment (Cao and Cheng, 2016; Uziela, et al., 2016). For instance, secondary structure prediction was widely utilized in the template-based structure modeling through threading or comparative modeling on those proteins that have structurally determined homologs (Roy, et al., 2010; Wang, et al., 2010; Webb and Sali, 2014), and in *ab initio* modeling for those proteins whose sequences share few sequential similarities with known solved structures (Kryshtafovych, et al., 2017; Ovchinnikov, et al., 2017).

The progress in protein secondary structure prediction over the past few decades can be generally summarized from two aspects: the discovery of novel features that are useful for prediction and the development of effective machine learning algorithms (Rost, 2001; Yang, et al., 2016). The early attempts utilized statistical propensities of single amino acid observed from known structures to identify secondary structures in proteins (Chou and Fasman, 1974). The subsequent improvements came from the inclusion of sequence evolutionary profile features inferred from multiple sequence alignment (MSA) such as position-specific scoring matrices (PSSM) (Altschul, et al., 1997; Dor and Zhou, 2007; Jones, 1999; Magnan and Baldi, 2014; Pollastri and Mclysaght, 2004; Pollastri, et al., 2002). In addition to the PSSM, the Hidden Markov model (HMM) profiles derived from HHblits (Remmert, et al., 2012) was proposed for predicting protein structural properties (Meng, et al., 2018). Atchley’s factors were also included in some studies to capture the similarity between the types of amino acids (Atchley, et al., 2005; Spencer, et al., 2015).

Meanwhile, the machine learning algorithms for protein secondary structure prediction also continued to improve. Several early approaches applied shallow neural networks (Holley and Karplus, 1989; Qian and Sejnowski, 1988), information theory and Bayesian analysis (Gibrat, et al., 1987; Schmidler, et al., 2000; Stolorz, et al., 1992) to secondary structure prediction. PSIPRED (Jones, 1999) method proposed a two-stage neural network to predict the secondary structure from the PSI-BLAST sequence profiles. SSpro (Pollastri, et al., 2002) used bi-directional recurrent neural networks to capture the long-range interactions between amino acids. Deep learning techniques recently achieved significant success in secondary structure prediction (Dor and Zhou, 2007; Fang, et al., 2018; Faraggi, et al., 2012; Heffernan, et al., 2017; Spencer, et al., 2015; Wang, et al., 2016). DNSS (Spencer, et al., 2015) applied an ensemble of deep belief networks to predict 3-state secondary structure. SPIDER2 (Heffernan, et al., 2015) employed stacked sparse auto-encoder neural networks to predict the several structural properties iteratively, and this method was further advanced by bidirectional long- and short-term memory (LSTM) neural networks to capture the long-range interactions (Heffernan, et al., 2017). DeepCNF (Wang, et al., 2016) integrated the convolutional neural networks with conditional random-field to learn the complex sequence-structure relationship and interdependence between sequence and secondary structure. Porter 5.0 (Torrisi, et al., 2018) ensembled seven bidirectional recurrent neural networks to improve the protein structure prediction. Assisted with the power of deep learning, the accuracy of 3-state secondary structure prediction has been successfully improved above 84% (Fang, et al., 2018; Heffernan, et al., 2017; Wang, et al., 2016) on some benchmark datasets.

In this work, we developed an improved version of our *ab initio* secondary structure method using multiple advanced deep learning architectures (DNSS2). Three major improvements have been made over the original DNSS method. Firstly, besides the PSSM profile features and Atchley’s factors used in DNSS, we incorporated several novel features such as the emission and transition probabilities derived from Hidden Markov model (HMM) profile (Remmert, et al., 2012), and profile probabilities inferred from multiple sequence alignment (MSA) (Magnan and Baldi, 2014). All the three new features represent the evolutionary conservation information for amino acids in sequence. Secondly, we designed and integrated six types of advanced one-dimensional deep networks for protein secondary structure prediction, including traditional convolutional neural network (CNN) (Krizhevsky, et al., 2012), recurrent convolutional neural network (RCNN) (Liang and Hu, 2015), residual neural network (ResNet) (He, et al., 2016), convolutional residual memory networks (CRMN) (Moniz and Pal, 2016), fractal networks (Larsson, et al., 2016), and Inception network (Szegedy, et al., 2015). The ensemble of six networks from DNSS2 significantly improved the secondary structure prediction. Finally, DNSS2 was trained on a large dataset, including 4,872 non-redundant protein structures with less than 25% pairwise sequence identity and 2.5 Å resolution. Our method was extensively tested on the independent dataset and the latest CASP13 dataset with other state-of-art methods and delivered the state-of-the-art performance.

## 2 Materials and Methods

### 2.1 Experimental design

In this work, the main objective was to improve the secondary structure prediction by developing more advanced deep learning architectures and introducing more useful features. In the process, we have developed a systematic framework to effectively build deep learning architectures and obtain features to improve secondary structure prediction. **Figure 1** provides an overview of our experimental design. **Figure 1(A)** lists the six major steps of designing, training and testing deep learning architectures. **Figure 1(B)** illustrates the process of creating training and validation datasets. The key analysis is to design appropriate architectures and investigate if they can improve prediction accuracy. Six different deep neural network architectures were evaluated in the study, including convolutional neural network (CNN) (Krizhevsky, et al., 2012), recurrent convolutional neural network (RCNN) (Liang and Hu, 2015), ResNet (He, et al., 2016), convolutional recurrent memory network (CRMN) (Moniz and Pal, 2016), FractalNet (Larsson, et al., 2016), and Inception network (Szegedy, et al., 2015). Most of these architectures were applied to secondary structure prediction for the first time. The detailed description of each network is included in Section 2.4. To ensure a fair comparison, each network was optimized using the original feature profiles of training proteins and evaluated on the same validation set of DNSS1. The network that achieved the best Q3 accuracy was selected to explore the feature space on the profiles derived from multiple sequence alignments (MSA) generated by PSI-BLAST (Altschul, et al., 1997) and HHblits (Remmert, et al., 2012), Atchley factors, and emission/transition probabilities inferred from the Hidden Markov model (HMM) profile. The optimal feature set was determined according to the highest Q3 accuracy on the validation datasets. The networks were then re-trained using the optimal input profiles to obtain the best models.

**Fig. 1.**
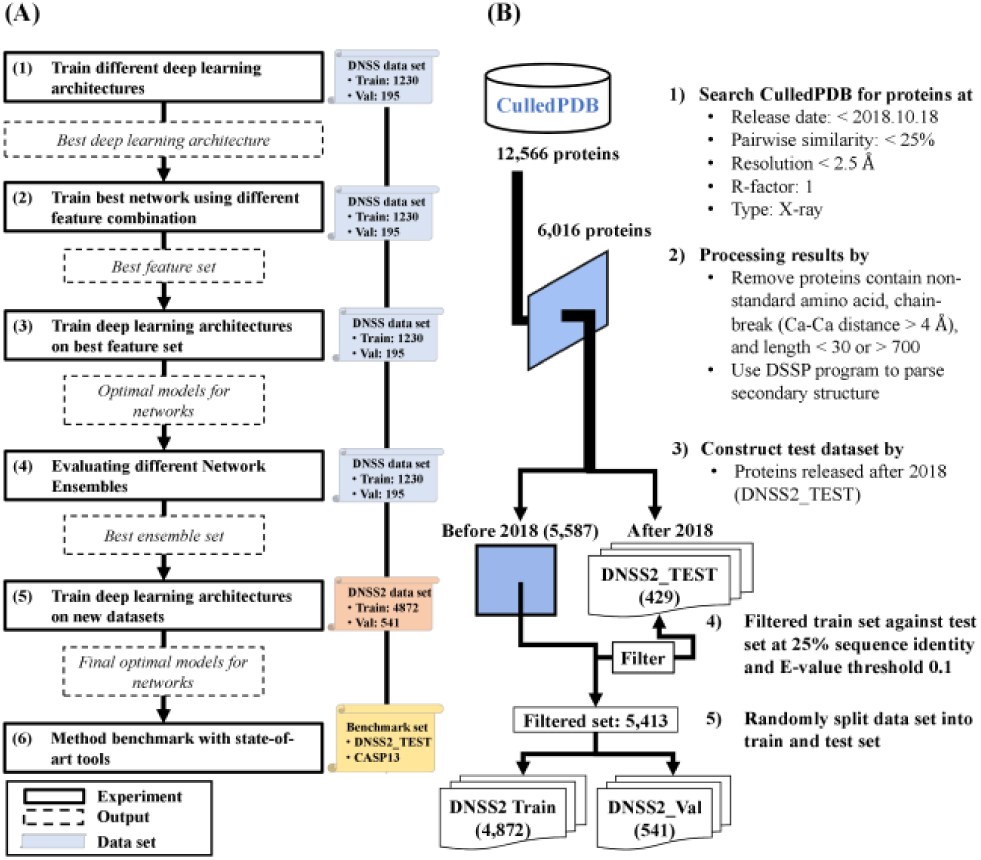
Overview of the experimental workflow for improving secondary structure prediction. (A) Six principal steps are conducted to construct and train deep networks. The solid box represents an analysis step. The dashed box represents the output from the previous step. The scroll represents the dataset used in each step. (B) Dataset generation and filtering process.

Since combining predictors generally improved the prediction accuracy, the different combinations of networks were also evaluated. Finally, after the optimal sets of deep learning architectures and feature profiles were determined, all networks were re-trained on the large dataset that was manually curated including the non-redundant proteins whose structures have been released publicly before 2018. The final networks were used to predict the secondary structure for the test proteins. The probabilities of the three states (i.e. helix, sheet, and coil) for each residue predicted by six networks were averaged to make the final secondary structure prediction.

Our method was then benchmarked with other state-of-art methods on the two independent test datasets.

### 2.2 Datasets and evaluation metric

As described in section 2.1, two training datasets were used in our experiment. In the first stage, the original DNSS dataset (Spencer, et al., 2015) that included 1,230 training proteins and 195 validation proteins was utilized to investigate whether the deep learning architectures and novel features can boost the prediction accuracy.

To utilize more data available since DNSS1 was published, a new, larger training set of DNSS2 was constructed from CullPDB (Wang and Dunbrack Jr, 2003) curated on 18 October 2018 (**Figure 1(B)**). The dataset consists of 12,566 proteins that share less than 25% sequence identity with 2.5 Å resolution cutoff and R-factor cutoff 1. The structures of all the proteins were determined by X-ray crystallography. The dataset was then filtered by removing proteins with non-standard amino acids, chain-break (i.e. distance of adjacent Ca-Ca atoms is larger than 4 Å), and sequence length shorter than 30 or longer than 700 amino acids. Considering all external methods benchmarked in this work were developed prior to year 2018, the proteins that were released after Jan 1^st^, 2018 were extracted as independent test set (DNSS2_TEST). The resulting set of proteins was further filtered against DNSS2_TEST set using CD-HIT suite (Li and Godzik, 2006) with criteria of 25% sequence identity cutoff and e-value threshold 0.1. Finally, 5,413 proteins released prior to Jan 1^st^, 2018 were obtained as our training set, in which 4,872 proteins were used for network training (DNSS2_TRAIN) and 547 proteins were used for model selection (DNSS2_VAL). In addition, the proteins of the CASP13 (2018) experiment were collected and the ones with at least 25% sequence identity with training proteins were removed, which results in a set of 82 test proteins. The proteins were also classified into template-based (TBM) and free-modeling (FM) targets based on the official CASP definition (CASP 13, 2018, http://www.predictioncenter.org/casp13/index.cgi). In summary, the final test set contain 429 proteins from DNSS2_TEST and 82 proteins from CASP13.

We evaluated our secondary structure prediction based on two primary metrics: Q3 accuracy and Segment Overlap measure (SOV). Q3 score represents the percent of correctly predicted secondary structure states in a protein. SOV score measures the similarity between the predicted segments of continuous structure states and those in the experimental structure (Spencer, et al., 2015; Zemla, et al., 1999). The Q3 and SOV scores are complementary with each other for secondary structure evaluation. All training and testing proteins’ structure files were parsed by DSSP program (Kabsch and Sander, 1983) to obtain the real secondary structure classification for each amino acid for training and evaluation.

### 2.3 Input features

The profile of each amino acid is represented by 21 numbers from PSI-BLAST-based position specific scoring matrix (PSSM), 20 emission probabilities and 7 transition probabilities extracted from Hidden Markov Model (HMM) profile, 20 probabilities of standard amino acid calculated from the multiple sequence alignment (MSA) and 5 numbers derived from Atchley’s factor. These features (73 numbers in total) represent the evolutionary conservation and physicochemical properties for residues in a protein sequence.

PSI-BLAST was run to generate multiple sequence alignment and PSSM profile through searching a sequence against filtered UniProt sequence database at 90% sequence identity (UniRef90) (Consortium, 2014) with three iterations and an e-value cutoff 0.001 (‘-evalue .001 -inclusion_ethresh .002’). Less stringent threshold was used (‘-evalue 10 -inclusion_ethresh 10’) in case some proteins did not have homologous sequences returned. In a PSSM profile, each position is represented by 20 numbers related to the probabilities for 20 standard amino acids appearing at the position in the multiple sequence alignment. In addition, the sequence information in the second to the last column in PSI-BLAST profile is given for each residue.

HMM profile was generated by running three iteration of ‘HHblits’ against the uniclust30 database (version: October 2017) (Mirdita, et al., 2016). Two types of probabilities were associated with each residue in a HMM profile: emission probability and transition probability. Emission probability represents the probability of a given amino acid occurring at the position in the multiple sequence alignment. The transition probability represents the probability transiting from an alignment state (i.e. match, insertion, and deletion) to another. Similar to PSSM, the emission frequencies of the 20 standard amino acid for each residue were reported in the HMM profile, and the probabilities were calculated according to formula:

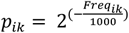

where *i* is the *i*-th residue in sequence and *k* is the *k*-th standard amino acid. And the probability is set to 0 if the frequency is denoted as ‘*’. The transition probabilities for each amino acid were also derived in the same fashion. In total, 20 emission probabilities and 7 transition probabilities for each amino acid were collected to represent the residue conservation inferred from HMM.

Since HHblits was more sensitive to identify distant homologous sequences than PSI-BLAST, the probability matrix of amino acids was also calculated from the multiple sequence alignment (MSA) generated by HHblits. The conversion from MSA to a probability matrix follows the same calculation as SSpro (Magnan and Baldi, 2014).

### 2.4 Deep learning architectures

A widely used deep learning architecture in bioinformatics is deep convolutional neural networks (CNN). Convolutional neural networks have some distinctive advantages over the traditional neural networks for the bioinformatics problems in several ways: (1) it can learn informative representation directly from sequence features without requiring segmentation (e.g. sliding window) or dimension reduction (e.g. principle component analysis) techniques; (2) the convolutional network can learn both local and global features to discover complex patterns; and (3) the architecture is independent of input size (i.e. length or volume). In this work, we design a standard CNN and five advanced deep learning architectures based on both convolutional and other useful operations as in **Figure 2.**

**Fig. 2.**
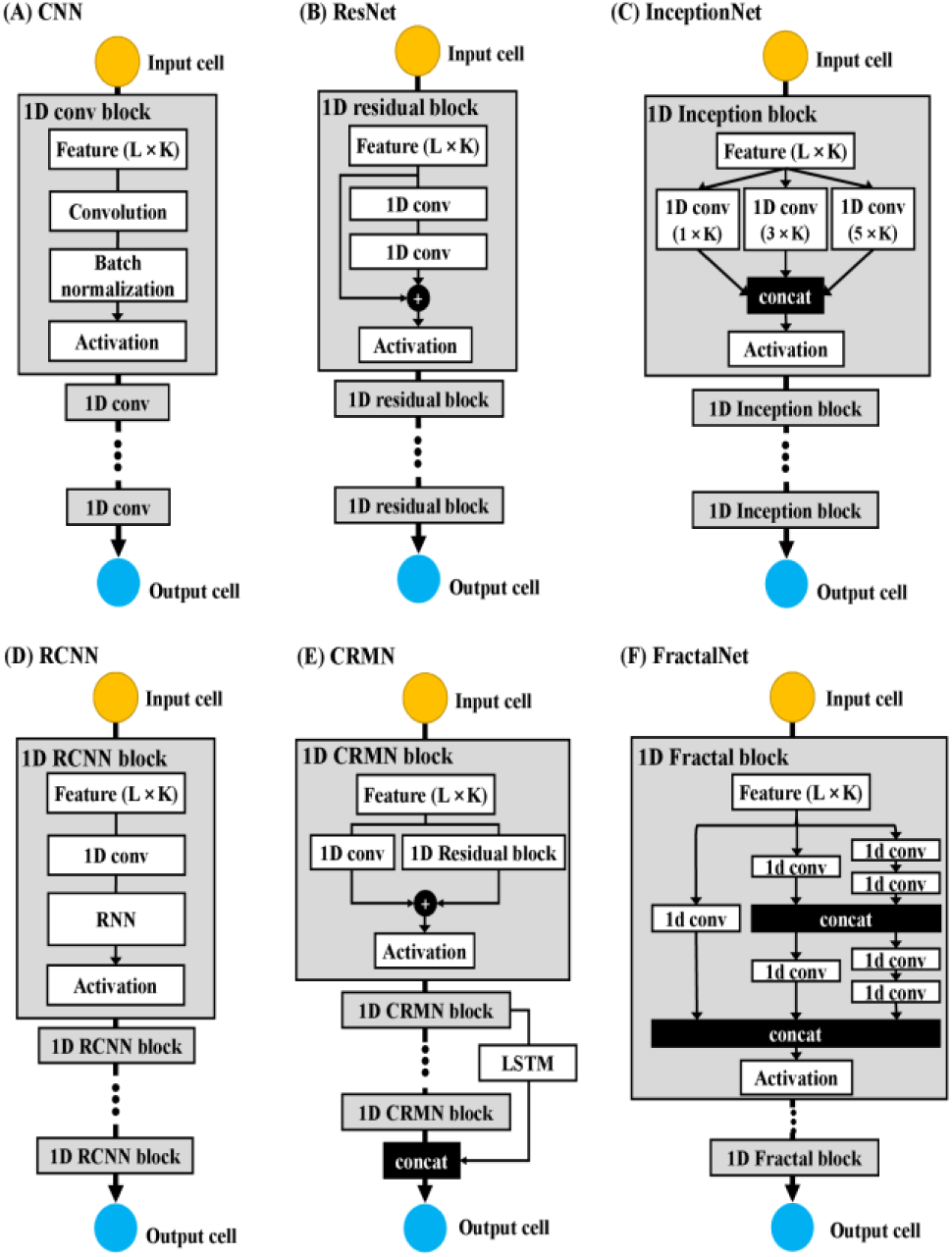
Six deep learning architectures: (A) CNN, (B) ResNet, (C) InceptionNet, (D) RCNN, (E) CRNN, (F) FractalNet) for secondary structure prediction. L: sequence length; K: number of features per position.

**Figure 2(A)** illustrates our standard convolutional neural network (CNN) for secondary structure prediction, consisting of a sequence of convolutional blocks, each of which contains a convolutional layer, a batch-normalization layer, and an activation layer. The original input is a L × K vector (*X*), where L is sequence length and K is the number of features per residue position in the sequence. For each convolution block, the feature maps are obtained after the convolution operation is applied by multiplying the weight matrices (called filters, *W*) with a window of local features on the previous input layer and adding bias vectors (*b*) according to the formula: *X*^*l*+1^ = *W*^*l*+1^ * *X*^*l*^ + *b*^*l*+1^, where *l* is the layer number. The batch normalization layer is added to obtain a Gaussian normalization of convolved features coming out of each convolutional layer. Then an activation function such as rectified linear function (i.e. ReLU) is applied to extract non-linear patterns of the normalized hidden features. To avoid overfitting, regularization approaches such as dropout (Srivastava, et al., 2014) can be applied in the hidden layers. The final output node (also a filter) in the output cell uses the softmax function to classify the input of each residue position from its previous layer into one of three secondary structure states. The output is a L × 3 vector, holding the predicted probability of three secondary structure states for each of L positions in a sequence. The final optimal CNN architecture includes 6 convolutional blocks, in which the filter size (window size) for each convolutional layer is 6, and the number of filters (feature maps) in each convolution layer is 40.

The residual network (ResNet) was designed to make traditional convolutional neural network deeper without gradient vanishing. The architecture constructs many residual blocks and stacked up them to form a deeper network, as shown in **Figure 2 (B)**. In each residual block, the input *X*^*l*^ is fed into a few convolutional layers to obtain the non-linear transformation output *G(X*^*l*+1^). In order to make the network deeper, an extra skip connection (i.e. short-cut) is added to copy the input *X*^*l*^ to the output of non-linear transformation layer, where *X*^(*l*+1)*^ can be represented as *X*^(*l*+1)*^ = *X*^*l*^ + *G(X*^*l*+1^) before applying another ReLU non-linearity. This process makes neural network deeper by adding shortcuts to facilitate gradient back-propagation during training and achieve better performance. The residual blocks with different configuration can be stacked to achieve higher accuracy. For instance, the final best architecture in DNSS2 is made up of 13 residual blocks, each of which includes 3 convolutional layers with filter size 1, 3, 1 respectively. The first three residual blocks used 37 filters to learn features, while the middle four blocks used 74 filters for each convolution layer, and the last six residual blocks used 148 filters. In total, 39 convolutional layers are included in the final residual network. In the network, the dropout and batch normalization were also added to prevent network from overfitting.

Inception network is an advanced architecture for building deeper networks by repeating a bunch of inception modules, as shown in **Figure 2(c)**. Instead of trying to determine the best values for certain hyper-parameters (i.e. number of filter size, number of layers, inclusion of pooling layer), inception network proposes to concatenate outputs of hidden layers with different configuration through an inception module and trains the network to learn patterns from the combination of diverse hyper-parameters. Despite its high computation cost, inception network has performed remarkably well in many applications (Fang, et al., 2018; Szegedy, et al., 2015). For secondary structure prediction, a combination of three filter sizes 1 × K, 3 × K and 5 × K was applied to convolve feature input, where K is the number of original input features for each residue position. The concatenation of the convolution outputs is fed into an activation layer for non-linear activation calculation. This kind of inception module is repeated to make a deeper network. After the parameter tuning, the optimal inception network is comprised of three inception blocks with 24 convolution layers included.

In addition, we designed three more deep learning architectures: recurrent convolutional neural network (RCNN) (Liang and Hu, 2015), convolutional residual memory networks (CRMN) (Moniz and Pal, 2016), and fractal network for secondary structure prediction. The recurrent convolutional neural network (RCNN) was designed to model sequential dependency hidden inside the sequential features (**Figure 2(D))**, It firstly extracts the higher-level feature maps by a convolution block, and then uses a recurrent neural network (i.e. bi-directional Long-Short-Term Memory (LSTM) network) for modeling the inter-dependence among the convolved features. Such a recurrent convolutional block with 4 convolutional layers included is repeated 5 times to build a deep recurrent convolutional neural network for secondary structure prediction in this work. The CRMN network augmented the architectures by integrating convolutional residual networks with LSTM (**Figure 2(E))** (e.g., 2 residual blocks and 2 LSTM in the network). Both methods advanced the convolutional neural network by introducing the memory mechanisms of recurrent neural network (RNN). Moreover, inspired by ResNet and Inception Network, we built a Fractal network stacking up different number of convolution blocks in both parallel and hierarchical fashion by adding several shortcut paths to connect lower-level layers and higher-level layers, as shown in **Figure 2(F)**. After tuning, the fractal network was assembled with 16 convolution layers for one fractal block.

### 2.5 Training and evaluation procedure

Deeper networks with complex architectures are generally difficult to train effectively due to the high-dimensional hyper-parameter space. To obtain good performance on specific feature sets within a reasonable amount of time for each deep network, we developed an efficient heuristic random sampling approach for model hyperparameter optimization. Specifically, based on the several trials on network training, we first determined heuristically a reasonable range for each type of the network hyperparameters, including the number of filters from 20 to 50, the number of convolution blocks from 3 to 7, and the filter size from 3 to 7. For each subsequent trial, the values of hyper-parameters were randomly sampled from their specified range and the Q3 accuracy of the network on the validation dataset under the specific parameter combination was assessed. For each deep network, the best parameter set was determined after 100 trials were evaluated. We found that using the random sampling technique was able to generate better models in most cases and was also more efficient than the traditional grid search or greedy search.

The performance of different deep architectures and different feature profiles on the secondary structure prediction were rigorously examined using the training and validation set from original DNSS method. After the parameters and input features were determined, we trained each deep network on the latest curated dataset (DNSS2_TRAIN) and selected best models using the Q3 accuracy on the independent validation dataset (DNSS2_VAL). We used the Keras library (http://keras.io/) along with Tensorflow as a backend to train all networks.

The performance of DNSS2 was evaluated on the two independent datasets and compared with a variety of the state-of-art secondary structure prediction tools, including SSpro5.2 (Magnan and Baldi, 2014), PSSpred (Yan, et al., 2013), MUFOLD-SS (Fang, et al., 2018), DeepCNF (Wang, et al., 2016), PSIPRED (McGuffin, et al., 2000), SPIDER3 (Heffernan, et al., 2017), Porter 5 (Torrisi, et al., 2018) and our previous method DNSS1 (Spencer, et al., 2015). All the methods were assessed according to the Q3 and SOV scores on each dataset.

## 3 Results

### 3.1 Benchmarking different deep architectures of DNSS2 with DNSS1

The first evaluation was to investigate whether the new deep architectures networks (DNSS2) outperform the deep belief network (DNSS1) for the secondary structure prediction. In order to fairly compare them, we trained and validated the six deep networks on the original input features of the same 1,230 training and 195 validation proteins used to train and test DNSS1. **Table 1** compares the Q3 and Sov scores of DNSS1 and DNSS2 architectures on the validation set. The results show that five out of six new advanced deep networks (RCNN, ResNet, CRMN, FractalNet, and InceptionNet) except the standard CNN network obtain higher Q3 scores than the deep belief network that used in DNSS1. InceptionNet worked best among individual deep architectures. The ensemble of the six deep architectures (DNSS2) achieved the highest Q3 score of 83.04%, better than all the six individual deep architectures and 79.1% Q3 score of DNSS1.

**Table 1.** Performance of the six different deep architectures and their ensemble on the DNSS1 validation dataset. DNSS2 represents the ensemble of six deep architectures (CNN, RCNN, ResNet, CRMN, FractalNet and InceptionNet).

| Method | Q3(%) | Sov(%) |
| --- | --- | --- |
| DNSS1 | 79.1 | 72.38 |
| DNSS2_CNN | 77.86 | 68.42 |
| DNSS2_RCNN | 79.87 | 72.34 |
| DNSS2_ResNet | 79.61 | 69.94 |
| DNSS2_CRMN | 79.32 | 69.21 |
| DNSS2_FractalNet | 79.85 | 72.82 |
| DNSS2_InceptionNet | 80.68 | 72.74 |
| DNSS2 | 83.04 | 72.74 |

### 3.2 Impact of different input features

After the best deep learning architecture (i.e. InceptionNet) was determined, it was utilized to examine the impact of the different input features including PSSM, Atchley factor (FAC), Emission probabilities (Em), Transition probabilities (Tr), and amino acids probabilities from HHblits alignments (HHblitsMSA). In this analysis, the protein sequence databases required for alignment generation were updated to latest and all the input features for DNSS1 datasets were regenerated. Specifically, the Uniref90 database that was released at October 2018 was used to generate PSSM profiles by PSI-BLAST, and the latest version of Uniclust30 database (October 2017) was used to generate HMM profiles by HHblits. The Inception network was then trained on the 1,230 proteins using the combination of five kinds of features. We tested six feature combinations shown in **Table 2**. Hyper-parameter optimization was applied to obtain the best model on each feature combination. **Table 2** shows the performance of different input feature combinations with the inception network on the validation dataset of 195 proteins. Adding the emission profile inferred from HMM model on top of PSSM and Atchley factor features increased the Q3 score from 79.81% to 82.31%. Integrating all the five kinds of features will yield the highest Q3 score (i.e. 82.72%) and Sov score (75.89%).

**Table 2.** Performance of different input feature combinations on the validation dataset of 195 proteins. PSSM, FAC, Em, Tr, HHblitsMSA denote five kinds of features: PSSM, Atchley factor, Emission probabilities, Transition probabilities, amino acid probabilities from HHblits alignments.

| Rank | Feature Name | Q3(%) | SOV(%) |
| --- | --- | --- | --- |
| 1 | PSSM + FAC + Em + Tr + HHblitsMSA | 82.72 | 75.89 |
| 2 | PSSM + FAC + Em + Tr | 82.36 | 76.03 |
| 3 | PSSM + FAC + Em | 82.31 | 74.15 |
| 4 | PSSM + FAC + HHblitMSA | 81.98 | 74.67 |
| 5 | PSSM + FAC + Tr | 80.13 | 71.61 |
| 6 | PSSM + FAC | 79.81 | 71.43 |

The performance of the six deep architectures and their ensemble on the latest features (the combination of all five kinds of features) of the DNSS1 validation dataset was also reported in **Table 3**. All six architectures were re-trained on the 1,230 proteins and evaluated on the validation dataset. Compared to the results in **Table 1**, the prediction accuracy of all the networks on the validation set was improved. The Q3 and SOV scores of the ensemble (DNSS2) were increased to 83.84% and 75.5%, respectively. The results indicate that the update of the protein sequence databases helps improve prediction accuracy.

**Table 3.** Performance of the six different deep learning architectures (CNN, RCNN, ResNet, CRMN, FractalNet, and InceptionNet) and their ensemble (DNSS2) on DNSS1 validation dataset and the updated protein sequence database.

| Method | Q3(%) | Sov(%) |
| --- | --- | --- |
| DNSS2_CNN | 80.29 | 72.1 |
| DNSS2_RCNN | 81.83 | 73.97 |
| DNSS2_ResNet | 81.53 | 73.71 |
| DNSS2_CRMN | 81.91 | 73.37 |
| DNSS2_FractalNet | 82.02 | 73.8 |
| DNSS2_InceptionNet | 82.74 | 75.3 |
| DNSS2 | 83.84 | 75.5 |

### 3.3 Comparison of DNSS2 with eight state-of-the-art tools on two independent test datasets

DNSS2 was compared with eight state-of-art methods including SSPro5.2, DNSS1, PSSpred, MUFOLD-SS, DeepCNF, PSIPRED, SPIDER3, and Porter 5 on the DNSS2_TEST dataset. The test dataset contains non-redundant proteins released after Jan 1^st^, 2018. All the tools were downloaded and configured based on their instructions. The sequence databases that the tools require were updated to the latest version.

The Q3 score of each tool on the test dataset was reported in **Table 4**. In general, DNSS2 is comparable to the two predictors (Porter 5 and SPIDER3) on this dataset and outperforms the other six methods. Specifically, DNSS2 achieved a Q3 accuracy of 85.02% and SOV accuracy of 76.01% on the DNSS2_TEST dataset, which was significantly better than DNSS 1.0 on the DNSS2_test dataset with p-value equal to 2.2E-16.

**Table 4.**
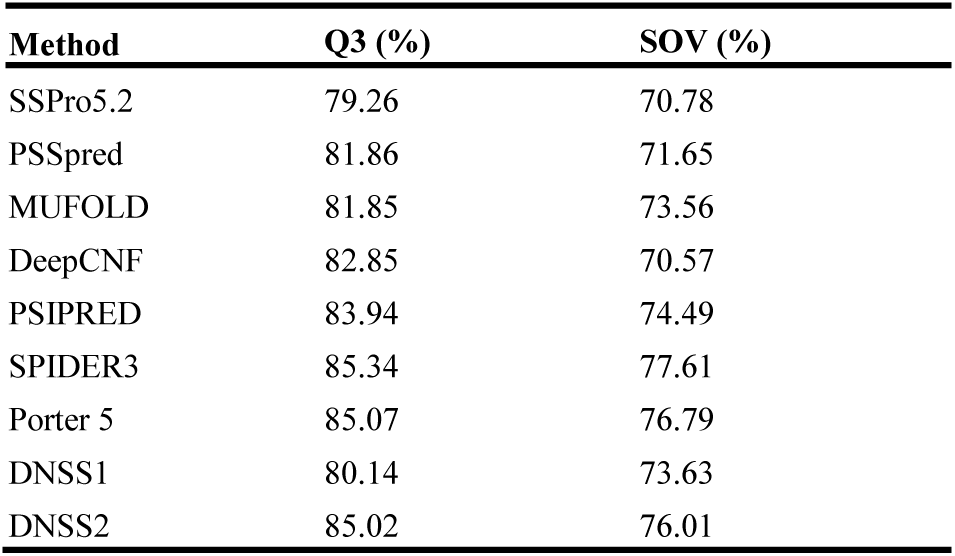
Q3 scores of 9 secondary structure prediction methods on DNSS2_test dataset. Three methods (SPIDER3, Porter5, DNSS2) have Q3 score higher than 85%.

| Method | Q3 (%) | SOV (%) |
| --- | --- | --- |
| SSPro5.2 | 79.26 | 70.78 |
| PSSpred | 81.86 | 71.65 |
| MUFOLD | 81.85 | 73.56 |
| DeepCNF | 82.85 | 70.57 |
| PSIPRED | 83.94 | 74.49 |
| SPIDER3 | 85.34 | 77.61 |
| Porter 5 | 85.07 | 76.79 |
| DNSS1 | 80.14 | 73.63 |
| DNSS2 | 85.02 | 76.01 |

In addition to the DNSS2_test dataset, we also compared these methods on the 82 protein targets of 2018 CASP13 experiment, which share less than 25% sequence identity with the training proteins of DNSS2. Both template-based (TBM) and free-modeling (FM) protein targets were used to evaluate the methods and the results are summarized in the **Table 5**. Consistent with the performance on the DNSS2_test dataset shown in **Table 4**, DNSS2, SPIDER3 and Porter 5 performed best, while DNSS2 achieved slightly better performance than SPIDER3 and Porter 5. **Figure 3** plots the distribution of the Q3 scores for all CASP13 targets obtained by DNSS2 and the other eight methods. In general, the distribution of DNSS2 consistently shifts to higher Q3 score compared with other methods, even though the distribution of DNSS2 largely overlaps with that of SPIDER3 and Porter 5.

**Table 5.**
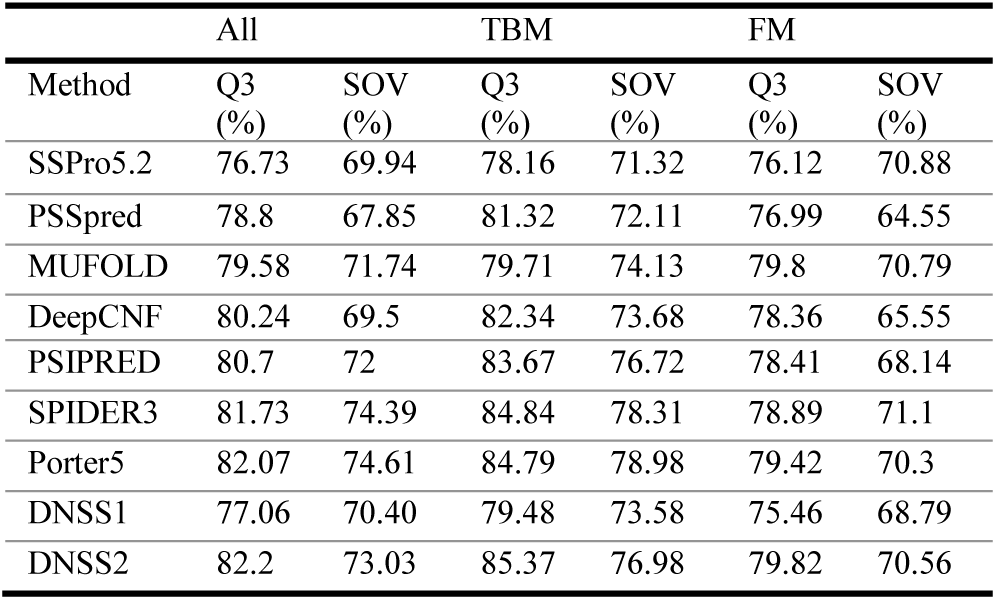
Comparison of methods on the CASP13 dataset in terms of all CASP13 targets, template-based targets, and template-free targets.

**Fig. 3.**
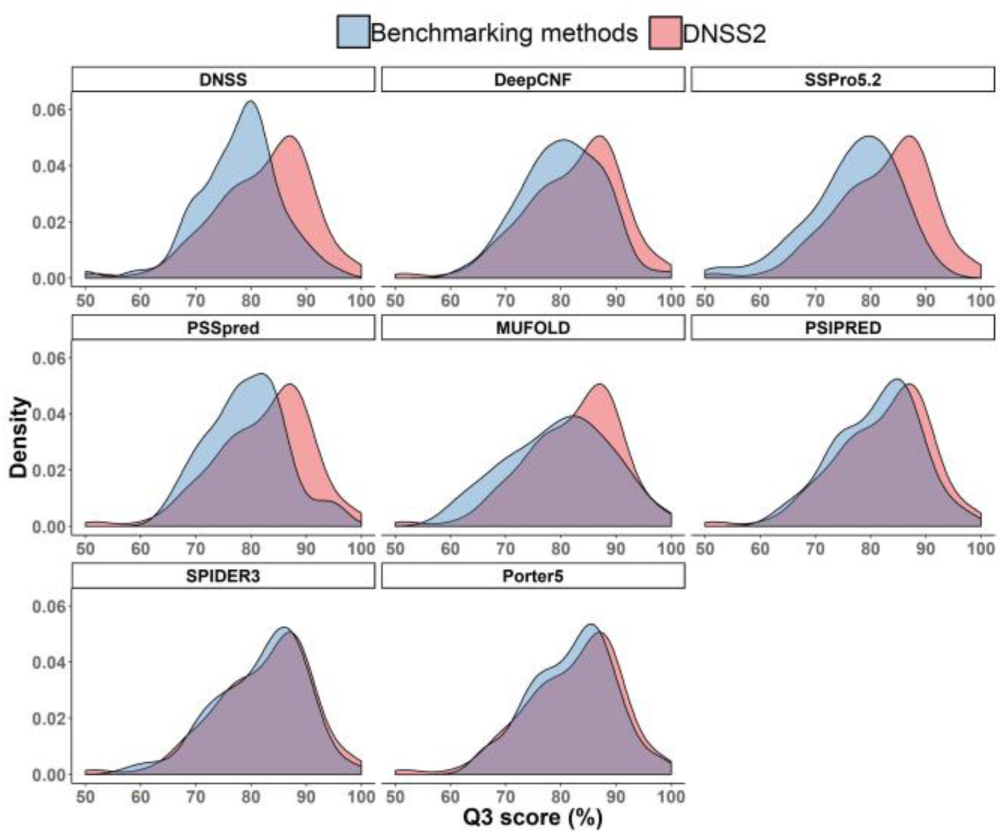
Comparison of the distribution of Q3 scores of eight existing methods and that of DNSS2 on all CASP13 targets.

**Table 6** summarized the confusion matrix of predictions of three kinds of secondary structures (helix, sheet, coil) by DNSS2 on the CASP13 dataset. DNSS2 yields the highest accuracy for helical prediction (87.91%), followed by the coil prediction (80.21%) and the sheet prediction (76.45%). The prediction errors between helix, sheet, and coil was also reported. The error rate of misclassifying helix as sheet is the lowest (0.57%) and sheet as coil is the highest (22.46%).

**Table 6.** Confusion matrix of helix, sheet and coil predicted by DNSS2 on CASP13 dataset.

|  | C pred | E pred | H pred |
| --- | --- | --- | --- |
| Coil (C) | 80.21% | 9.51% | 10.28% |
| Sheet (E) | 22.46% | 76.45% | 1.10% |
| Helix (H) | 11.52% | 0.57% | 87.91% |

## 4 Conclusion

In this work, we developed several advanced deep learning architectures and their ensemble to improve secondary structure prediction. We investigated six advanced deep learning architectures and five kinds of input features on secondary structure prediction. Several deep learning architectures such as inception network, fractal network, and recurrent convolutional memory network are novel for protein secondary structure prediction and performed better than the deep belief network. The performance of the deep learning method is comparable to or better than seven external state-of-the-art methods on the two independent test datasets. Our experiment also demonstrated that emission/transition probabilities extracted from hidden Markov model profiles are useful for secondary structure prediction.

